# The Secreted Metabolite Isopentenyladenine from *Faecalibacterium prausnitzii* Mitigates Gut Inflammation

**DOI:** 10.1101/2025.08.24.671981

**Authors:** Lina Yao, Angelo Solania, Anny-Claude Luissint, Aaron T. Balana, Hua Zhang, Dewakar Sangaraju, Zijuan Lai, James Kuo, Kelly M Storek, Dennis W. Wolan

## Abstract

Colonic microbiome dysbiosis is correlated with inflammatory bowel disease (IBD), and depletion of the commensal bacterium *Faecalibacterium prausnitzii* (*F. prausnitzii*) is routinely observed in the metagenomic analyses of IBD patient microbiome samples. *F. prausnitzii* is likely beneficial to hosts, as oral administration of *F. prausnitzii* strain A2-165 has anti-inflammatory properties in murine models of colitis. Previous studies attribute the anti-inflammatory effects of *F. prausnitzii* A2-165 to production of the short-chain fatty acid butyrate, as well as a 15 kDa protein known as microbial anti-inflammatory molecule (MAM). Here, we verified that oral dosing of strain A2-165 protects against DSS-induced murine colitis and further show the aqueous-soluble secreted fraction of overnight cultures from a collection of *F. prausnitzii* strain inhibit inflammatory signatures, including the activation of the host’s NF-κB pathway, production of IL-8, and differentiation of naïve T cells into the T_H_17 lineage. We find that MAM is secreted in extracellular vesicles; however, MAM-containing vesicles do not have anti-inflammatory properties in our collection of assays and suggests that MAM is likely not a direct contributor. Untargeted and targeted mass spectrometry metabolomics analyses on the soluble anti-inflammatory secretome yielded several unique *F. prausnitzii* metabolites, including isopentenyladenine. We demonstrated isopentenyladenine independently modulates host cellular signaling and immune responses and suggests this newly identified metabolite with human immunomodulatory properties may be useful towards the discovery of IBD-focused therapeutics.

## Introduction

The human gut microbiome is critical for maintaining human health^1^. This complex microbial ecosystem is essential for preserving intestinal barrier integrity, modulating immune responses, and ensuring gut homeostasis. Emerging evidence suggests that microbiome dysbiosis, including loss of microbial diversity and beneficial commensal organisms, is a hallmark of ulcerative colitis (UC) and Crohn’s disease (CD). *F. prausnitzii* is consistently one of the most diminished species in IBD, with its reduction closely linked to disease severity, poor surgical outcomes, and higher recurrence rates following surgery^2–4^. In line with these observations, recent data from Genentech’s GARDENIA phase 3 clinical trial^5^ further support its potential relevance in treatment response. These findings reinforce the relevance of *F. prausnitzii* as a potential biomarker in biologic therapy response and highlight the complexity of host-microbe interactions in the IBD setting.

*F. prausniztii* is a butyrate-producing commensal bacterium. The anti-inflammatory effects of butyrate have been well characterized, as the molecule enhances intestinal epithelial barrier integrity via upregulation of tight junction proteins^13^ and epithelial cell energy metabolism^14–16^. However, butyrate alone does not fully account for the broad immunomodulatory properties of *F. prausniztii*.

Recent reports suggest the presence of additional active factors that contribute to its therapeutic potential in gut inflammation. Broadly, administration of *F. prausnitzii* strain A2-165 to TNBS-, DNBS-, and DSS-induced colitis murine models has substantial protective benefits, including significant reduction in colonic inflammation, improvement in mucosal barrier integrity, and suppression of pro-inflammatory cytokine expression such as TNF-α, IL-6, and IL-17A^3,6–8^. Additionally, *F. prausnitzii* administration promotes the production of anti-inflammatory molecules such as IL-10, modulates the gut microbiota composition by enhancing beneficial microbial populations, and reduces oxidative stress in the gut environment^3,9–11^. A2-165 has also been shown to attenuate weight loss and improve histological scores in experimental colitis models, further underscoring its potential as a therapeutic strategy for IBD^3,8,12^.

In addition to butyrate, other studies have suggested a primary beneficial constituent secreted by *F. prausnitzii* includes a protein termed microbial anti-inflammatory molecule (MAM)^3^ that influences the inhibition of the NF-κB pathway and suppression of pro-inflammatory cytokine production by unknown mechanisms^17^. We find that MAM is secreted in extracellular vesicles (EVs) and MAM-containing EVs did not yield anti-inflammatory responses that were observed with the *F. prausnitzii* supernatant and suggest that MAM is not a primary functional mediator of *F. prausnitzii*’s immunomodulatory effects in our collection of assays. Notwithstanding our result, *F. prausnitzii*-derived EVs may play a role in mediating its immunomodulatory effects, although the precise mechanisms are still under investigation^15,18^. The full potential repertoire of host-beneficial factors produced by *F. prausnitzii* and the mechanism(s) of action by which the bacterium exerts its immunomodulatory functions require further investigation.

T cells, including the T_H_17 differentiated lineage, are pivotal to promoting inflammatory immune responses within the intestinal mucosa and are implicated in IBD pathogenesis ^19,20^. T_H_17-driven inflammation is mediated by the secretion of cytokines, such as IL-17A that promotes chronic intestinal inflammation. As such, targeting T_H_17 differentiation and reducing the production of pro-inflammatory molecules is a promising therapeutic strategy in IBD, as well as other inflammatory diseases. In line with previous studies, we demonstrate that cultured *F. prausnitzii* supernatants suppress T_H_17 cell differentiation, modulate systemic inflammation *in vivo*, and inhibit the NF-κB signaling pathway.

Microbiome metabolomics studies have increasingly highlighted the critical role of microbial metabolites in promoting host health, particularly through anti-inflammatory effects and the enhancement of barrier function, such as butyrate, regulates immune responses by modulating T_reg_ cell differentiation and suppressing pro-inflammatory cytokines^21,22^. Other microbial metabolites, such as tryptophan-derived indole compounds, influence the aryl hydrocarbon receptor (AhR) pathway, reducing intestinal inflammation and improving barrier function^23^. Additionally, bile acid derivatives metabolized by gut microbes have been shown to exert anti-inflammatory effects by modulating signaling pathways such as FXR, TGR5, and RORγt^20,24^. These findings underscore the need to discover and characterize microbial metabolites as such molecules represent potential leads towards identifying novel therapeutic strategies to manage inflammatory diseases and maintain gut health.

The immune-modulatory effects of metabolites from gut microbes are increasingly recognized; however, few examples link the biologically active molecules with mechanisms of action responsible for the observed effects. Our search for additional *F. prausnitizii* anti-inflammatory factors yielded isopentenyladenine (iP), an adenine-based cytokinin typically associated with plant signaling, as a previously unrecognized immunomodulatory compound in the human gut microbiota context. Using targeted metabolomics and functional assays, we demonstrated that iP significantly suppresses NF-κB activation, inhibits T_H_17 cell differentiation, and promotes epithelial barrier integrity, revealing a potent, multi-pronged anti-inflammatory effect. iP and butyrate likely act in concert to modulate host signaling pathways. These findings broaden our understanding of host-microbiome interactions and open new avenues for the development of microbiome-based therapies for IBD and other inflammation-driven diseases.

## Results and Discussion

### Live *F. prausnitzii* Protects Against DSS-induced Colitis in Mice

We investigated the extent of protection *F. prausnitzii* A2-165 provides mice in DSS-induced colitis. Treatment groups received daily oral administration of 1 × 10⁹ CFU live bacteria, purified cultured supernatant, heat-inactivated bacterial culture supernatant, or vehicle control (*e.g.,* YCFA media) for eleven days. DSS was introduced in the drinking water to induce colitis two days after initiation of bacterial gavage and was provided for seven consecutive days. Mice were weighed daily, and disease severity was assessed two days post DSS exposure (day 9) by measuring colon length and secreted cytokine serum levels of individual mice.

While previous studies have demonstrated that *F. prausnitzii* A2-165 confers protective effects in colitis models, we extended these findings by incorporating a broader panel of readouts, including systemic cytokine profiling, correlation analyses, and comparison across live and inactivated bacteria. Mice receiving live *F. prausnitzii* A2-165 showed significantly improved body weight recovery post-DSS exposure compared to colitis control mice, while no improvement was observed in groups treated with culture supernatants or heat-inactivated bacteria (**Fig. 1a**).

**Fig. 1.**
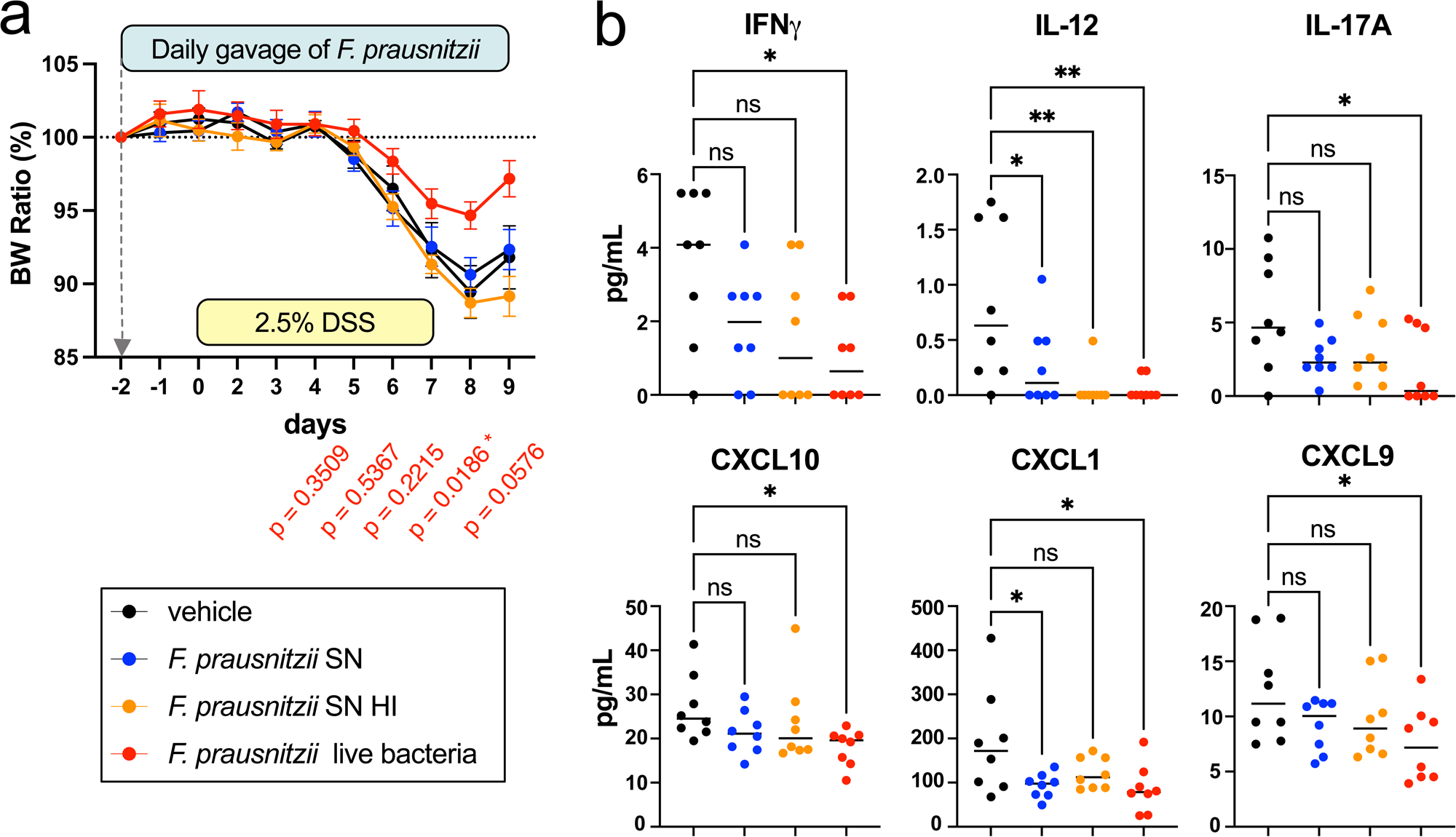
| Live *F. prausnitzii* protects in the DSS-colitis mouse model. Briefly, mice were orally gavaged daily with one of the following: vehicle control (culture media YCFA with glycerol), *F. prausnitzii* supernatant (SN), heat-inactivated *F. prausnitzii* supernatant (SN HI), or *F. prausnitzii* live bacteria (1x10^9^ in 200 µL) from Day -2 to Day 8. Daily body weight changes of mice. A significant difference in body weight was observed between the live *F. prausnitzii* treatment group and the vehicle group on Day 8 (**a**). Cytokine levels in mouse sera collected at experimental endpoint from the DSS-colitis experiment, as measured by Luminex in triplicate. Significantly altered cytokines include IFN-γ, IL-12, IL-17, CXCL10, CXCL1, and CXCL9 (**b**). Data points below the limit of detection were treated as zero. The full cytokine panel is shown in **Fig. S1**. Data are presented as mean ± SEM. Statistical significance was determined using one-way ANOVA with Dunnett’s multiple comparisons.

Serum samples collected on day 9 were analyzed using a Luminex Assay Panel. Several inflammatory cytokines and chemokines, including IL-17A, IFN-γ, IP-10, CXCL1, and CXCL9 were reduced in all treatment groups, significance was only observed with live bacteria gavage relative to the colitis control (**Figs. 1b, S1**). IL-6, a key cytokine upstream of T_H_17 responses, was also measured. Notably, we observed inverse correlations between IL-17A and IL-6 serum levels and body weight on days 8 and 9 body in colitic mice treated with live *F. prausnitzii* (**Figs. S2a, S2b**), and these cytokines were also inversely correlated with colon length at the endpoint (**Figs. S2c, S2d**). These findings not only confirm previous reports of *F. prausnitzii*’s protective role in colitis, but also provide new mechanistic insight into its systemic immunomodulatory effects.

### *F. prausnitzii* Supernatants Suppress T_H_17 Cell Differentiation

IL-17A is a hallmark of T_H_17-mediated inflammation and reduced serum levels of IL-17A have been consistently reported in studies exploring the anti-inflammatory properties of the gut microbiome^26,27^. IL-17A-producing T_H_17 cells contribute to chronic inflammation through the recruitment of neutrophils and the promotion of pro-inflammatory cytokines. Therefore, the observed cytokine shifts in this study, particularly the reductions in IL-17A and the correlations with clinical markers of disease severity, highlight a potential immunomodulatory role of *F. prausnitzii* via the T_H_17 axis. Given the established role of the T_H_17 pathway in intestinal inflammation and autoimmune diseases, our results prompted further investigation into how *F. prausnitzii* modulates T_H_17-driven immune responses.

We explored the direct regulation of *F. prausnitzii* strains isolated from a collection of human donors^28^ on the differentiation of naïve murine T cells to mature T_H_17 T cells. Supernatants derived from *F. prausnitzii* isolates grown in YCFA medium were incubated with naïve T cells isolated from wild-type C57BL/6J (B6Jax) mice under T_H_17-cell differentiation conditions. All *F. prausnitzii* isolate supernatants significantly inhibited the differentiation of naïve CD4+ T cells from wild-type C57BL/6J mice into T_H_17 cells (**Fig. 2a**) with no or minimal impact on cell viability (**Fig. S3a**), except for two isolates that exhibited poor growth. Our analyses also suggest that the extent of reduction in T_H_17 cells is strongly correlated with density of the cultured bacteria, as those *F. prausnitzii* isolates with higher culture densities (as measured by OD_600_) were more effective at preventing T cell differentiation (**Fig. 2b**). Given that greater *F. prausnitzii* cell numbers were associated with enhanced suppression of T_H_17 differentiation, this raises the possibility that a secreted factor or metabolite may mediate this effect.

**Fig. 2.**
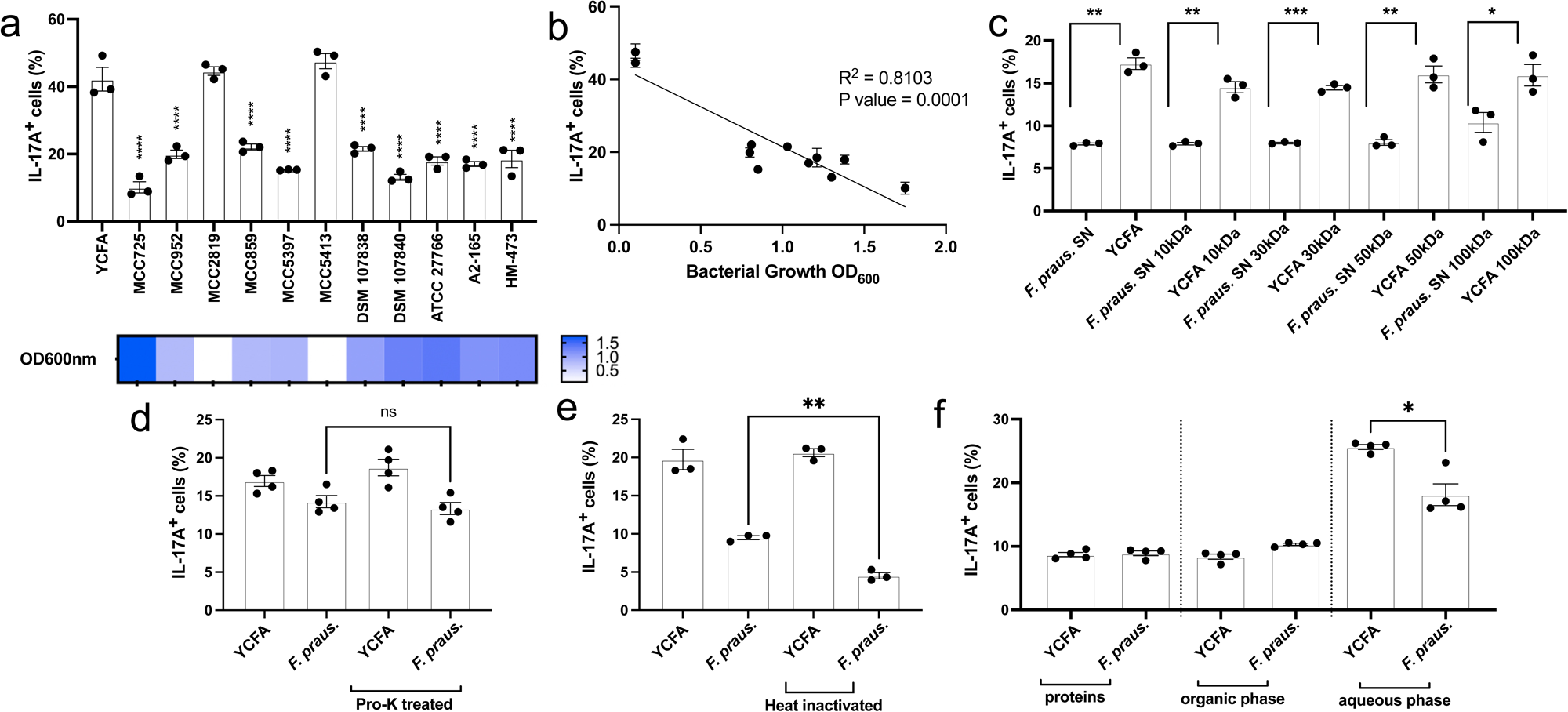
| *F. prausnitzii* culture supernatants inhibit T_H_17 differentiation. Supernatants from a representative of *F. prausnitzii* strains inhibited murine naïve T cell differentiation to T_H_17 with population frequencies of T cells activated and expanded *in vitro* (**a**). The inhibition level of T_H_17 cells directly correlates with the growth density of the *F. prausnitzii* cultures (**b**). All fractions, including the whole supernatants and filtrates obtained by 10 kDa, 30 kDa, 50 kDa, and 100 kDa MWCO filters from the *F. prausnitzii* A2-165 culture, exhibited significant inhibition of T_H_17 cell differentiation in vitro when compared to their corresponding components of the YCFA medium controls (**c**). Notably, the proteinase K-treated supernatants (**d**) from the *F. prausnitzii* A2-165 culture retained their ability to inhibit T_H_17 cell differentiation, whereas heat-inactivated supernatant (**e**) showed a further reduction in this activity, suggesting the involvement of temperature-stable components. Only the aqueous phase fraction of *F. prausnitzii* A2-165 culture supernatants demonstrated significant inhibition of T_H_17 cell differentiation in vitro (**f**). The values are expressed as the mean ± SEM in percentage of IL-17A^+^. n = 3 biologically independent samples per group. One-way ANOVA was used following the YCFA medium control column as the follow-up tests. TEM images of *F. prausnitzii* A2-165 samples, including a medium control (YCFA), the original (1x) culture supernatant treated with proteinase K (PK) and heat inactivation, 10x concentrated supernatant treated with PK and heat inactivation, and 10x concentrated supernatant treated with PK and PMSF inactivation. Top panel shows images at 45,000× magnification, and bottom panel at 1,200,000× magnification, using negative staining (**g**). Purified EVs show no inhibitory effect on T_H_17 differentiation across the tested concentration range (**h**).

We subjected A2-165 supernatants to size-based fractionation to isolate and identify the anti-inflammatory factor(s). Whole supernatant and filtrates obtained through 10 kDa, 30 kDa, 50 kDa, and 100 kDa molecular weight cut-off (MWCO) filters all significantly reduced T_H_17 cell differentiation compared to their respective YCFA culture medium-treated controls (**Figs. 2c, S3b**), suggesting the active product(s) are likely small molecules and/or peptides smaller than 10 kDa. Whole supernatants were treated with proteinase K (Pro-K) to assess the role of proteinaceous components in the T_H_17 inhibitory activity. Welch’s t-test showed no significant difference between the Pro-K-treated and untreated supernatants, further suggesting that intact proteins are unlikely to be the primary mediators of the effect (**Figs. 2d, S3c**). Notably, when the supernatant was subjected to heat treatment, a significant reduction in activity was observed (**Figs. 2e, S3d**), indicating that temperature-sensitive impurities may have been removed, thereby contributing to the enhanced effects observed. To further explore the biochemical nature of the active compounds, A2-165 supernatants underwent organic solvent extraction with chloroform-methanol. Only the aqueous-soluble fraction (top layer) retained its inhibitory effect on T_H_17 differentiation (**Figs. 2f, S3e**), while the organic extraction phase (bottom layer) and protein-isolated fraction (white interphase layer) showed no activity. Our findings suggest that *F. prausnitzii* A2-165 produces small, heat-stable, and Pro-K-resistant metabolites capable of suppressing T_H_17 cell differentiation *in vitro*, and the identification of the secreted factors could help guide the development of therapeutics against T_H_17-mediated inflammation.

### *F. prausnitzii* Secretes MAM in Extracellular Vesicles

Given these observations, we next sought to determine whether the T_H_17-inhibitory phenotype was mediated by MAM, as MAM has reported anti-inflammatory properties. We generated MAM-specific polyclonal antibodies to detect if MAM is produced in A2-165 overnight culture supernatants. Western blot analyses showed MAM is one of the most prominently secreted proteins and was notably resistant to Pro-K treatment (**Fig. 3a**). These observations combined with the fractionation results (*i.e.,* MWCO filtrates and aqueous fraction (**Figs. 2c, 2f**)), prompted further investigation into the role of MAM as a stable protein whether involved in other immune modulation.

**Fig. 3.**
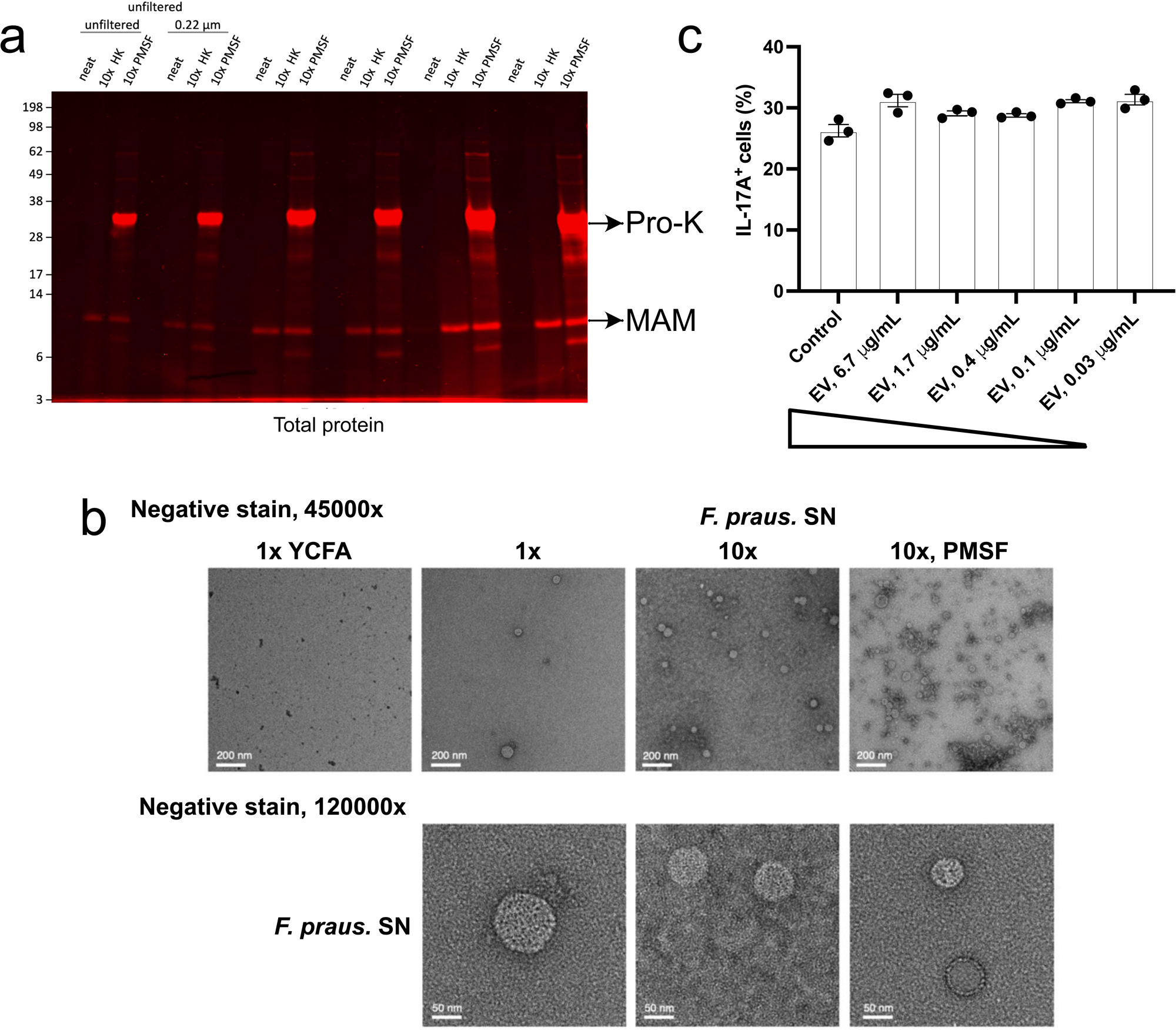
| *F. prausnitzii* A2-165 MAM is secreted in EVs. TEM images of *F. prausnitzii* A2-165 samples, including a medium control (YCFA), the original (1x) culture supernatant treated with proteinase K (PK) and heat inactivation, 10x concentrated supernatant treated with PK and heat inactivation, and 10x concentrated supernatant treated with PK and PMSF inactivation. Top panel shows images at 45,000× magnification, and bottom panel at 1,200,000× magnification, using negative staining (**a**). Samples used for microscopy in the negative staining (**b**).

Supernatants were analyzed by transmission electron microscopy (TEM) to establish the localization (*e.g.,* surface-bound and/or full secretion) of MAM. As previously reported, TEM analysis showed *F. prausnitzii* produces and secretes 50-100 nm extracellular vesicles (EVs) and are within the range of typical bacterial EV sizes (**Fig. 3b**). Native staining reveals the total protein content of each sample and fraction, including those containing MAM within these vesicles, as well as within the lysate pellet (**Figs. S4a, S4b**). MAM-containing EVs were isolated by ultracentrifugation and assessed in the murine T_H_17 differentiation assay, and we observed no inhibition of T cell differentiation in a dose-dependent assay (**Figs. 3c, S3f**). Homology analysis of the MAM protein in strain A2-165 across the tested *F. prausnitzii* isolates revealed that strain MCC859 lacks the MAM homolog; however, MCC859 has T_H_17-inhibitory activity (**Fig. 2a**). These findings suggest that the reported anti-inflammatory role of MAM is independent of T_H_17 production and may modulate the immune response indirectly.

### *F. prausnitzii* Supernatants Suppress IL-8 Secretion and NF-κB Activation in Human Cells

DSS-induced colitis is not a T_H_17-driven model and prior reports investigating the host responses to *F. prausnitzii* demonstrated that supernatants significantly inhibited IL-1β-induced secretion of IL-8 in Caco-2 human epithelial cells^29,30^. *F. prausnitzii* supernatants also upregulate anti-inflammatory cytokines, such as IL-10 and TGF-β1^3,31^, and implicate the bacteria to have broader immunomodulatory functions and anti-inflammatory potential. IL-8 remains a key chemokine involved in neutrophil recruitment and epithelial inflammation. Given that *F. prausnitzii* has been reported to suppress IL-8 production in intestinal epithelial cells, we sought to explore whether this suppression may contribute to its broader anti-inflammatory effects. This is particularly relevant in the context of epithelial responses during intestinal injury. Therefore, we assessed the effects of *F. prausnitzii* A2-165 supernatant fractions on IL-8 secretion in HT29-MTX human colonic epithelial cells stimulated with TNFα. Treatments included overnight culture supernatant, <10 kDa MWCO flow-through, EV-depleted supernatant, graded doses of purified EVs, and sterile YCFA medium as a control. Bacterial supernatants were effective at significantly reducing IL8 levels (**Fig. S5a**); however, isolated EVs did not result in a cellular response or reduction of IL8 secretion (**Fig. S5b**).

We next tested if *F. prausnitzii* supernatants regulate NF-κB activity. Supernatants from all cultured *F. prausnitzii* isolates in our panel revealed inhibitory properties against NF-κB activation in a reporter assay (**Figs. 4a, S3g**). Supernatant fractions and isolated EVs from strains A2-165 were further investigated. Isolated EVs showed no dose-dependent effect on NF-κB activity in comparison to its vehicle control (PBS) (**Figs. 4b, S3h**). However, whole supernatants and the aqueous phase extracts from strains A2-165 and MCC859 suppressed NF-κB activity, and the active constituents were present in MWCO 10 kDa filtrates and unaffected by Pro-K exposure (**Figs. 4c, 4d**).

**Fig. 4.**
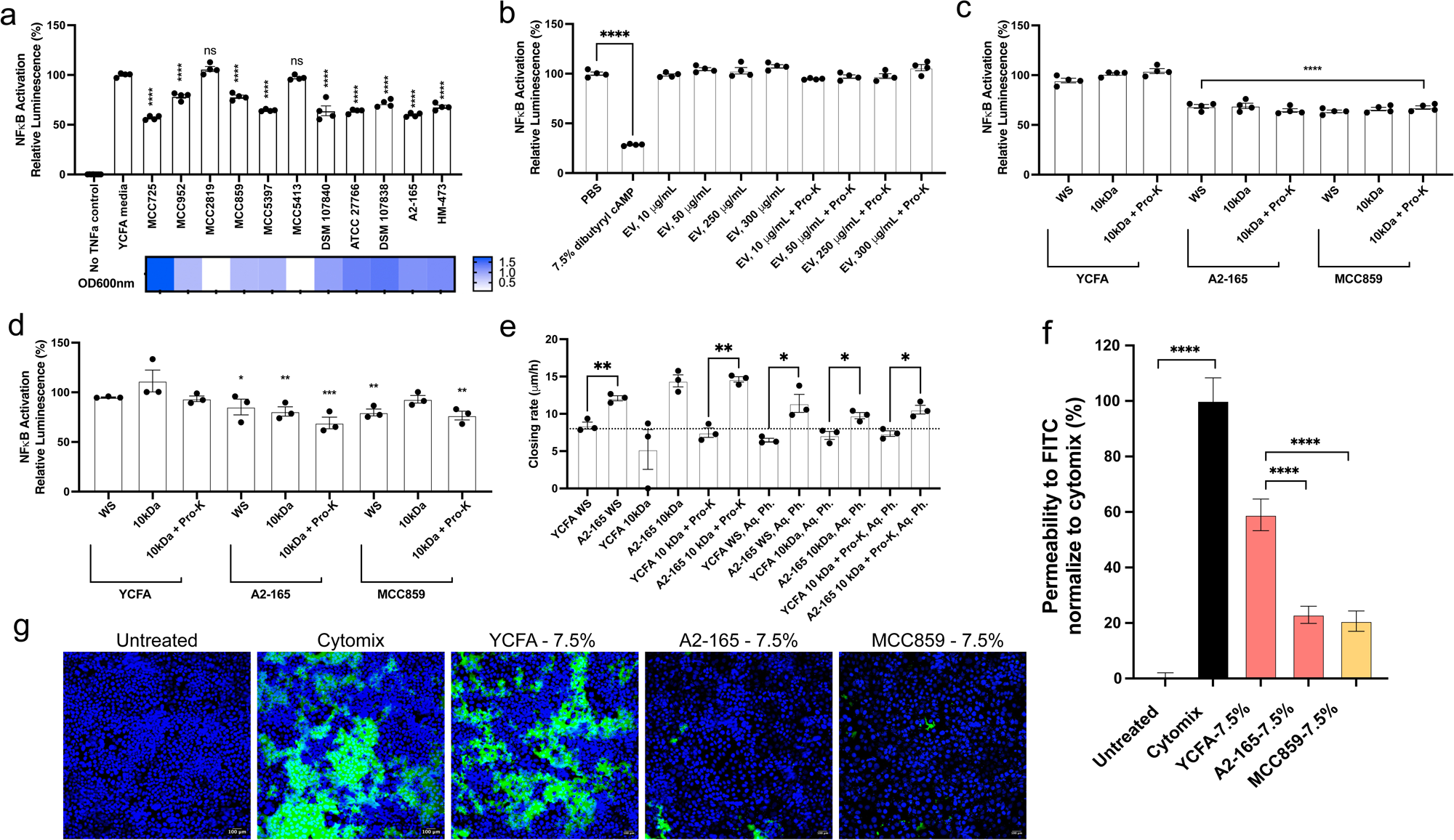
| *F. prausnitzii* culture supernatants suppress NF-κB activation and protect epithelial barriers. All *F. prausnitzii* strains tested showed a significant suppression of NF-κB activation using their supernatants, with only two strains exhibiting low growth density exceptions (**a**), while the extracted EVs from *F. prausnitzii* A2-165 did not exhibit such an effect (**b**). Dibutyryl cAMP works as a positive control which inhibits NF-κB activation. Whole supernatant, 10 kDa filtrate, and 10 kDa-fitrate with PK treated samples from both *F. prausnitzii* isolates displayed NF-κB suppression activation compared to their relevant controls (**c**). NF-κB suppression activity was exclusively observed in the aqueous phases from both A2-165 and MCC850 isolates (**d**). TNFα was added in the tested samples unless otherwise stated. The whole supernatant, the <10 kDa fraction followed by PK treatment of the supernatant, and the extracted aqueous phases of the *F. prausnitzii* A2-165 all demonstrated the promotion of cell differentiation/ proliferation as depicted in an epithelial wound closure assay using T84 human colonic epithelial cells (**e**). Supernatants from both *F. prausnitzii* strains, A2-165 and MCC859, demonstrated significant barrier-protective effects in an XPerT permeability assay using Caco-2 cells. Histograms display the mean fluorescence intensity of NeutrAvidin-DL488 across 3 replicates, normalized to the cytomix-treated condition, set as 100% leakage (**f**). Representative images show Caco-2 monolayers, selected from 8 images captured per well. Each condition was performed in triplicate. Green fluorescence depicts NeutrAvidin-DL488 (leakage), while blue fluorescence represents the cell nuclei. Scale bar: 100 µm. (**g**). One-way ANOVA was used following the corresponding fraction of the YCFA medium control column as follow-up tests.

Given that DSS disrupts the gut barrier, *F. prausnitzii*’s protective effects may also involve gut barrier restoration. We subjected *F. prausnitzii* to a scratch assay with T84 human colonic epithelial cells to evaluate barrier recovery ^32^. Remarkably, the addition of *F. prausnitzii* A2-165 whole supernatant, 10 kDa-filtered fraction or the Pro-K-treated supernatant accelerated scratch closure rates (**Fig. 4e**). Using a modified xPerT pertpermeability assay^33^, in which a proinflammatory cytokine cocktail (TNF-α, IFN-γ, IL-6, and IL-1β) was applied to mimic the inflammatory environment of the gut and disrupt epithelial barrier integrity, both *F. prausnitzii* strains A2-165 and MCC859 exhibited robust barrier-protective effects. These strains effectively countered the epithelial integrity disruption induced by cytokine challenge when compared to YCFA media alone. (**Figs. 4f**, **4g**).

### Small Molecule Metabolites Secreted by *F. prausnitzii* Drive Are Responsible for Host Effects

*F. prausnitzii* A2-165 and MCC859 supernatants, aqueous phase extracts, and YCFA medium were assessed by mass spectrometry-based untargeted metabolomics analyses to identify secreted metabolite(s) that may contribute to the observed effects. We identified nine small-molecule metabolites, including butyrate, with a >10-fold change relative to YCFA (**Table S1**). The samples were reanalyzed with targeted mass spectrometry to determine concentrations of each metabolite in the supernatants. We further confirmed the enrichment of several metabolites, including N-acetylhistamine, isopentenyladenine (iP), and butyrate, are produced by all *F. prausnitzii* samples and the abundance of these metabolites correlated with bacterial growth (**Fig. 5a**). Of note, N-acetylcadaverine and N-butyrylputrescine were produced in a select group of strains and nicotinic acid was the only detectable metabolite depleted across all *F. prausnitzii* samples compared to YCFA control (**Fig. 5a**).

**Fig. 5.**
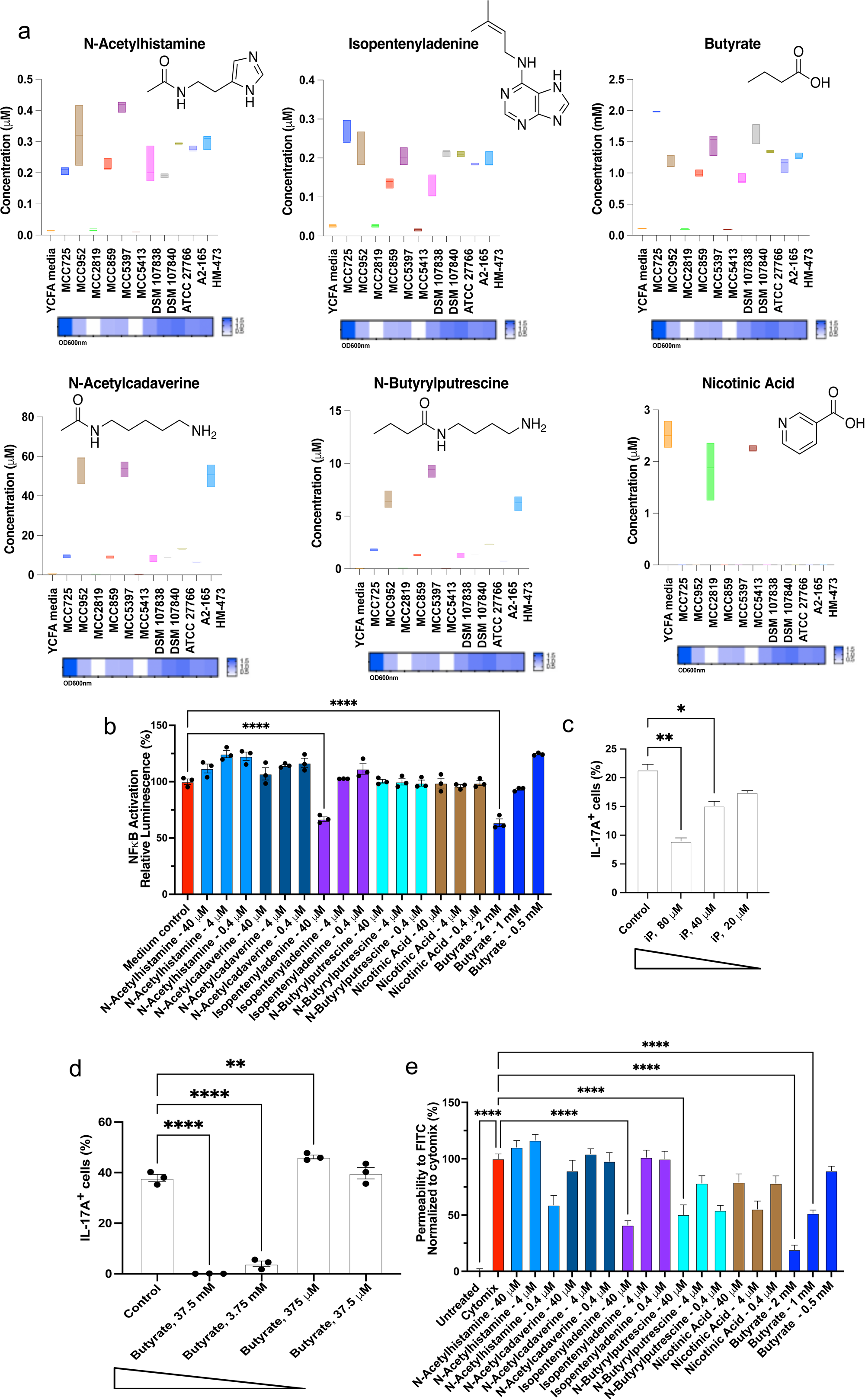
*F. prausnitzii* secreted isopentenyladenine and butyrate, which suppress NF-κB and inhibit T_H_17 cell differentiation. *F. prausnitzii* supernatants were subjected to untargeted and targeted metabolomics. N-acetylhistamine, N-acetylcadeverine, N-butyrylputrescine, isopentenyladenine (iP), and butyrate are notably enriched, while nicotinic acid is significantly depleted (**a**). Enrichment or depletion levels correlate with bacterial growth density, as shown in the heatmap below each graph. Both iP and butyrate suppress NF-κB activity (**b**). iP significantly suppresses T_H_17 differentiation across tested concentrations. 0.2% DMSO was used as the control (**c**). Sodium butyrate significantly suppresses T_H_17 differentiation in a dose-dependent manner at millimolar concentrations (left) but loses this effect at lower doses (right). 7.5% PBS was used as the control (**d**). iP and butyrate exhibit significant protective effects in an XPerT permeability assay using Caco-2 cells (**e**). Representative images display the barrier integrity of Caco-2 cell monolayers at the highest concentration of individual molecules. Green fluorescence depicts NeutrAvidin-DL488 (leakage), while blue fluorescence represents the cell nuclei. Scale bar: 100 µm (**f**). One-way ANOVA with Dunnett’s multiple comparisons was used for statistical analysis.

We investigated each enriched small molecule identified from our targeted MS analyses individually for NF-κB inhibition and found both butyrate and iP significantly inhibited NF-κB activation (**Fig. 5b**). Interestingly, iP and butyrate also exhibited dose-dependent T_H_17 inhibition (**Figs. 5c-5d, S3i-S3j**), suggesting a potential therapeutic relevance in inflammatory conditions. However, these effects were observed at supraphysiological levels, raising questions about their relevance in vivo, particularly during infection. In contrast, the activity of iP was detectable at much lower, more physiologically relevant concentrations (**Fig. 5a**).

To evaluate whether inhibition of inflammation translates to improved epithelial barrier function, we employed xPerT assay to measure restoration of paracellular barrier integrity in response to individual molecules. Remarkably, both iP and the well-characterized metabolite butyrate significantly neutralized the negative effects of cytokine-induced disruption on epithelial barrier integrity at higher concentrations (**Figs. 5e**, **6**), consistent with their inhibitory effects on NF-κB activation (**Fig. 5b**).

**Fig. 6.**
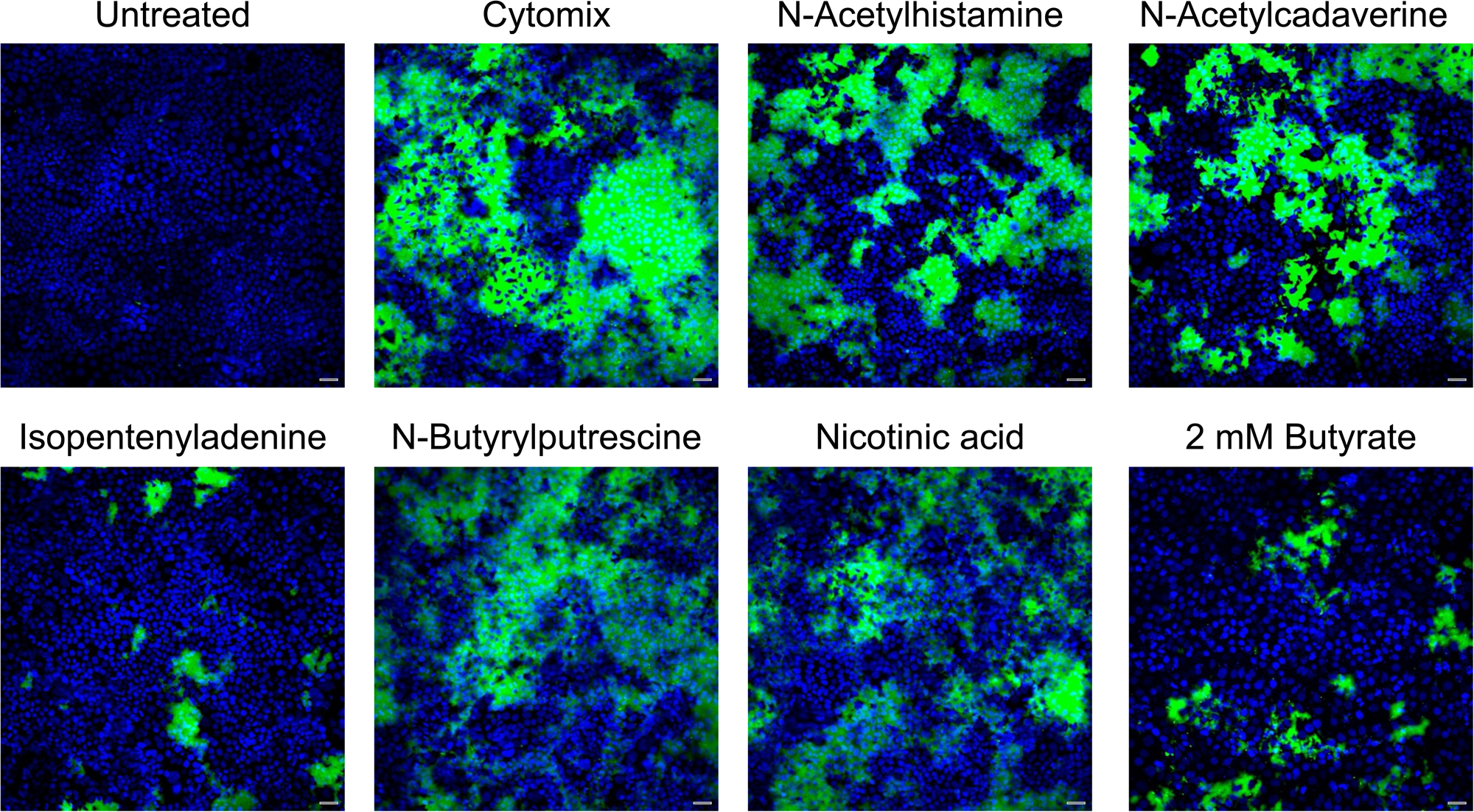
iP and butyrate preserve barrier integrity in an XPerT permeability assay using Caco-2 cells. Representative images display the barrier integrity of Caco-2 cell monolayers at the highest concentration of individual molecules. Green fluorescence depicts NeutrAvidin-DL488 (leakage), while blue fluorescence represents the cell nuclei. Scale bar: 100 µm.

Although the physiological concentrations of butyrate and iP in our assay were higher than those measured in the *F. prausnitzii* supernatants, butyrate levels in the human gut can reach 10-25 mM depending on diet, microbial composition, and health status^34–36^. The physiological gut levels of iP remains to be determined in both healthy individuals and IBD patients. Of note, live *F. prausnitzii* will produce elevated concentrations of these metabolites locally at the mucosal interface. Furthermore, a synergistic interplay among metabolites, including butyrate and isopentenyladenine, could contribute to the observed immunomodulatory and barrier protection effects.

Cytokinins, such as iP and the corresponding riboside derivatives, possess bioactivities beyond their canonical plant hormone functions^37^. iP is a weak inhibitor of cyclin-dependent kinases (CDKs)^38^ and the riboside conjugate isopentenyladenosine (iPR) protects against oxidative stress via Nrf2 pathway activation^39^. These molecules also have reported anti-inflammatory effects, including reduced neutrophil infiltration in mouse models of inflammation. iPR and other cytokinins are also agonists of the A3 adenosine receptor (A3R), known for its anti-inflammatory and antitumor activities^40^. Specifically, A3R activation by compounds like 2-Cl-IB-MECA have been shown to attenuate colitis and TNFα-induced inflammation through NF-κB signaling inhibition^41^. Given the structural and functional similarities between iP and its riboside derivatives, it is plausible that iP may also engage adenosine receptor pathways or other glucocorticoid receptor-mediated mechanisms. These connections warrant further investigation to elucidate the precise molecular targets of iP in epithelial barrier protection. Our findings underscore the potential of iP as a novel therapeutic candidate for diseases characterized by compromised barrier integrity and chronic inflammation, such as IBD.

## Conclusion

Our findings provide new key insights into the immunomodulatory properties and barrier protection effects of *F. prausnitzii* with the identification of the secreted anti-inflammatory metabolite iP. While we confirmed that culture supernatants from *F. prausnitzii* inhibit mouse T_H_17 differentiation while also suppressing IL-8 secretion, NF-κB activation, and enhancing barrier integrity in human cell lines. These outcomes are likely interconnected through a shared mechanism centered on NF-κB inhibition. As a master regulator of inflammation, NF-κB controls the transcription of numerous pro-inflammatory genes, including IL-8, and influences cytokine environments that drive T_H_17 polarization, such as IL-6 and IL-23. Therefore, dampening NF-κB signaling may simultaneously reduce epithelial-derived chemokine responses and interfere with the cytokine milieu required for T_H_17 differentiation. This integrated framework provides a unifying explanation for the broad anti-inflammatory effects observed across our in vitro and ex vivo assays. The collective activity of small molecules such as butyrate, iP, and potentially other yet-unidentified metabolites or peptides likely contribute to this coordinated immunoregulatory effect.

The physiological concentration of iP in the human gut, especially in health versus disease, remains to be established. Elucidating the host receptor(s) or signaling pathways through which iP exerts its effects will be critical to uncovering its mechanism of action. Finally, clarifying whether iP acts independently or synergistically with other microbial metabolites, such as butyrate, will be essential for understanding its full potential as a new avenue for therapeutic investigation. Addressing these knowledge gaps will guide the development of targeted microbiome-based interventions for IBD and other inflammatory diseases.

## Methods

### In Vivo DSS Mouse Experiment

Following, Eight-week-old C57BL/6J mice were purchased from Jackson Labs, acclimatized to our animal facilities, and assigned to treatment groups. Groups of mice received 200 μL of either live *F. prausnitzii* A2-165 bacterial culture (around 1 × 10⁹ CFU in 200 μL), bacterial culture supernatant (*e.g.,* depleted of bacteria), heat-inactivated supernatant, or sterile YCFA control medium (n = 8). Treatments were administered daily via intragastric gavage, beginning two days prior to the introduction of DSS. Colitis was induced by providing 2.5% (w/v) DSS in drinking water for seven days. Mice were monitored daily for body weight changes, clinical symptoms of colitis, and overall health. At the conclusion of the experiment (*e.g.,* day 9, **Fig. 1b**), mice were euthanized by cervical dislocation. Disease severity was assessed by measuring colon length, and quantitating levels of IL-17A, IFN-γ, IP-10, CXCL1, CXCL9, and IL-6 cytokines in sera using Luminex Assay Panels. Statistical analyses using one-way ANOVA with Dunnett’s multiple comparisons were performed to compare treatment groups and determine the protective efficacy of *F. prausnitzii* administration.

### *F. prausnitzii* Bacterial Culture

*F. prausnitzii* isolates were obtained from Microbiotica and were cultured in an anaerobic chamber (Coy Laboratory Products) under a gas mixture of 5% hydrogen, 20% carbon dioxide, and balance nitrogen. Individual strains were streaked from glycerol stocks onto YCFA agar and incubated for 48 h at 37 °C. Colonies were subsequently inoculated into 3 mL of YCFA liquid medium in Falcon™ round-bottom polystyrene tubes. Cultures were grown anaerobically for 48 h. Following incubation, bacterial cultures were harvested by centrifugation at 3,000 g for 10 min. Supernatants were collected by filtration through a 0.2 μm membrane filter to ensure sterility. These filtered supernatants were used in subsequent cell-based experiments, including T cell differentiation, IL8 secretion, and NFκB activity. All experiments were conducted in triplicate and repeated independently at least twice to ensure reproducibility and statistical validity.

### Untargeted and targeted metabolomics analysis for in vitro bacterial cultures

#### Untargeted metabolomics

All metabolite extraction procedures were kept on ice. Briefly, 350 μL of cold methanol containing in-house metabolomics Recovery IS (Stable Isotope Labeled Internal Standards) mixture and 200 μL of cold chloroform were added to each bacterial culture sample. Samples were vortexed for 5 minutes and centrifuged for 10 minutes. Next, supernatants were transferred and 200 μL cold H2O were added to perform liquid-liquid phase separation. Samples were mixed and centrifuged again. The top layer aliquots were transferred, dried, and reconstituted in 100 μL acetonitrile:water (8:2, v/v) containing the in-house metabolomics Global IS (Stable Isotope Labeled Internal Standards) mixture and submitted for global metabolomics analysis. Additionally, a derivatization assay targeting small organic acid metabolites was performed with details reported previously ^42^. For global metabolomics, ACQUITY UPLC BEH Amide column (2.1 mm × 150 mm × 1.7 μm, 130Å, Waters Corporation) was used to separate metabolites with mobile phase A of 100% H2O containing 10 mM ammonium formate and 0.125% FA, and mobile phase B of 95% acetonitrile in H2O containing 10 mM ammonium formate and 0.125% FA. For derivatized metabolites, Phenomenex Luna Omega C18 column (2.1 mm × 100 mm × 1.6 μm, 100Å, Phenomenex) was applied to chromatographically separate the analytes with mobile phase A of water containing 0.1% FA and mobile phase B of methanol only. Data acquisition was achieved on a Shimadzu Nexera HPLC series system (Shimadzu) coupled with a Thermo Q Exactive Plus Hybrid Quadrupole-Orbitrap Mass Spectrometer (Thermo Fisher Scientific). Injection volume of 3 μL was used for sample analysis under heated electrospray ionization (HESI) condition. Samples were run for both positive and negative modes. The Q Exactive Plus Mass Spectrometer was operated with the following parameters: sheath gas flow rate, 50 units; aux gas flow rate, 13 units; aux gas temperature, 425°C; capillary temperature, 263°C; spray voltage, 3500 V for pos and −2500 V for neg; scan mode, full MS scan with data-dependent MS/MS acquisition. In full MS scan, scan range is 60 to 900 m/z; resolution is 70,000; AGC target, 1×e^6; Maximum IT, 200 ms. In ddMS2 scan, the top five ions are selected with isolation window of 1.5 m/z; resolution is 17,500; AGC target, 5×e^4; Maximum IT, 20 ms. In-house metabolomics data processing software was used for metabolite identification and peak picking/peak integration. Metabolites were identified at Level 1 confidence ^43^ by matching at least two independent orthogonal experimental data (accurate mass, isotopic ratio, retention time, and MS/MS fragmentation pattern) against in-house compound library. Trend analysis of stable isotope labeled internal standards and matrix PoolQC samples were examined (with %RSD < 15%) to validate system suitability and data robustness. For each metabolite, relative quantification was obtained through either MS peak area of analyte or MS peak area ratio of analyte/internal standard.

#### Targeted metabolomics

Authentic standard compounds for metabolites of interest and corresponding stable isotope-labeled internal standards were obtained commercially. Calibration standard curve containing a mixture of all analytes were prepared at concentrations of 9.77, 19.53, 39.06, 78.13, 156.25, 312.50, 625.00, 1250.00, 2500.00, 5000.00 ng/mL by serial dilution. During batch analysis, 25 μL of sample aliquot (calibration standards and bacterial culture samples) with stable isotope-labeled internal standards spiked in at 100 ng/mL were processed. Detailed sample preparation procedures and LC-MS analysis were described above in the untargeted metabolomics method section. For data analysis, a small molecule compound database for metabolites of interest and corresponding stable isotope-labeled internal standards, including their retention times, chemical formulas, as well as accurate mass targeted ions, was constructed for compound identification. Then, peak detection and peak integration were achieved using the closest RT strategy and high-resolution targeted ions with 5 ppm mass accuracy. Finally, calibration curves and absolute quantification results were computed using linear regression model between analyte concentration and peak area ratio of analyte vs. corresponding IS.

### T_H_17 differentiation assays using mouse primary cells

Naive CD4⁺ T cells were isolated from spleens and lymph nodes of C57BL/6J mice (6–8 weeks old) followed by Naive CD4+ T Cell Isolation Kit (Miltenyi Biotec). Cells were seeded into plates pre-coated with hamster IgG and cultured in T cell medium (RPMI with 10% FBS, glutamine, 2-mercaptoethanol, penicillin-streptomycin (P/S)). T cell receptor activation was induced using anti-CD3 and anti-CD28 antibodies and T_H_17 differentiation was initiated by introduction of IL-6 and TGF-β1 as described^20,44^. Bacterial culture supernatants or small molecules were added 18 h post-TCR activation, and cells were harvested on day 3. Flow cytometry was used to analyze cytokine expression, using phorbol myristate acetate, ionomycin, and GolgiPlug stimulation prior to staining for surface and intracellular markers. Data acquisition and analysis were performed on BD flow cytometers using FlowJo software.

### IL-8 Human Cell Secretion Assay

The mucus-secreting human intestinal epithelial cell line HT29-MTX was cultured in complete Dulbecco’s Modified Eagle Medium (DMEM) supplemented with 10% fetal bovine serum (FBS) and 1% penicillin-streptomycin (P/S) at 37 °C in a 5% CO₂ atmosphere. Cells were seeded at a density of 4 × 10⁴ cells per well in 96-well plates and allowed to differentiate for approximately eight days until confluent monolayers formed. The culture medium was replaced every 2–3 days to maintain cell viability. On the day of the experiment, cells were washed with fresh DMEM containing 5% FBS. Test samples including fractions of *F. prausnitzii* supernatant, extracellular vesicles (EVs), or small molecules of interest were added simultaneously with 5 ng/mL of pro-inflammatory cytokine TNF-α to stimulate an inflammatory response. After 6 h of incubation, cell supernatants were harvested for analysis of IL-8 secretion. IL-8 concentrations were quantified using a DuoSet ELISA Development System (R&D Systems, DY208-5) following the manufacturer’s protocol. Assays were performed in triplicate to ensure reproducibility, and results were analyzed to assess the anti-inflammatory properties of *F. prausnitzii* and its metabolites on epithelial cell-mediated cytokine production.

### NF-κB Activation Assay Using NanoLuc Reporter Cell Line

HEK293 cells stably expressing the NF-κB-responsive NanoLuc reporter (InvivoGen) were cultured in complete DMEM supplemented with 10% FBS and and 1% P/S maintained in a 5% CO₂ incubator at 37 °C. Cells were seeded at a density of 3 × 10⁴ cells per well in white-walled 96-well plates and allowed to adhere overnight. On the day of the assay, cells were treated with fresh DMEM containing 5% FBS and 1% P/S. TNF-α (50 ng/mL) was used as the stimulus for NF-κB activation alongside experimental treatments with bacterial culture supernatant, extracellular vesicles (EVs), or purified small molecules. Final concentrations of test samples ranged from 0.4 μM to 40 μM, depending on prior dose-response analyses. Cells were incubated with treatments for 6 h. NanoLuc luciferase activity, reflecting NF-κB pathway activation, was measured using the Nano-Glo® Luciferase Assay System (Promega) according to the manufacturer’s instructions. Luminescence readings were recorded using a microplate luminometer, and data were normalized to untreated controls. All conditions were tested in triplicate, and experiments were repeated independently at least twice for reproducibility.

### Barrier Function Assay Using Scratch Assay and Xpert Image Assay

Human epithelial cell lines (T84 for scratch assay and Caco-2 for Xpert image assay) were cultured in complete DMEM supplemented with 10% FBS and 1% P/S at 37 °C in a humidified atmosphere containing 5% CO₂. Cells were seeded in 96-well plates at a density of 4 × 10^4^ cells per well and grown to confluence to form a monolayer. For barrier function assays, cells were pretreated with test compounds (*e.g*., *F. prausnitzii* supernatant or small molecules of interest) for 24 hours before conducting the scratch or introducing insults. Appropriate vehicle controls were used to ensure consistency and baseline comparisons.

The scratch assay was performed to assess the migratory capacity and barrier function of the epithelial cell monolayer. The Incucyte 96-well Woundmaker (Sartorius) was used to create a uniform scratch across the cell monolayer in each well. The medium was replaced with 0.25% FBS DMEM containing test samples at the indicated concentrations. Cells were incubated for an additional 12–48 hours, depending on the experimental design, and measured on a real-time imaging system (Incucyte). The rate of wound closure was calculated from the distance of the scratch area that had been closed at various time points, comparing the treated groups with the control group.

XPert permeability assay was conducted using black, optically clear flat-bottom 96-well PhenoPlates (50-209-9831, FischerScientific) based on the method described by Dubrovskyi et al. (2013) with minor modifications^33^. Gelatin from porcine skin (G 2500, Sigma) was biotinylated using EZ-Link NHS-LC-LC-Biotin (21343, Thermo Fisher Scientific) to achieve a final concentration of 10 mg/ml. 50 µl of the biotinylated gelatin solution was added to each well and allowed to adsorb overnight at 4°C. The wells were then washed twice with 200 µl of PBS without Ca and Mg. Caco-2 cells were seeded onto the biotinylated gelatin-coated plates at a density of (5 x 10^4^ cells per well) and cultured for 72 hours prior to testing. To disrupt epithelial barrier integrity, a pro-inflammatory cytokine cocktail containing 25 ng/ml TNF-α (210-TA-100/CF), 100 ng/ml IFN-γ (285-IF-100/CF), 25 ng/ml IL-6 (7270-IL-025/CF), and 20 ng/ml IL-1β (201-LB/CF) in Opti-MEM™ I Reduced-Serum Medium (31985062, Thermo Fisher Scientific) was applied 24 hours before the addition of bacterial supernatants or test compounds for an additional 24 hours. 1.25 mM sodium butyrate was used as a positive control. For imaging preparation, a solution containing 12.5 µg/ml NeutrAvidin-DL488 (22832, Thermo Fisher Scientific) and Hoechst (H21486, Thermo Fisher Scientific) was directly added to the culture medium and incubated for 30 minutes before terminating the experiment. Cells were fixed with 16% PFA for 30 minutes at room temperature, followed by three washes with 200 µl of PBS without Ca and Mg. Matrix-bound NeutrAvidin-DL488 (488 nm / 528 nm channel for leakage) and cell nuclei (392 nm / 440 nm, Hoechst) were imaged using the ImageXpress® Micro Confocal system (Molecular Devices) with 10 x objectives. For each condition, three wells were utilized, and eight images were captured per well. Data were analyzed as the mean fluorescence intensity of the 488 nm channel across replicates, normalized to the cytomix-treated condition, which was set as 100% leakage. An increased fluorescence intensity indicated higher permeability and compromised barrier function.

### Characterization of MAM-Containing Extracellular Vesicles (EVs) and Visualization EV Isolation and Visualization

*F. prausnitzii* MAM-containing EVs were purified from bacteria grown for 48 h, as described^45^. Cultures were verified as mid-log phase, which is optimal for EV production (Yaron et al., 2020). Bacterial cells were removed by centrifugation at 4,000 × g for 20 min at 4 °C and the supernatant containing the EVs was carefully collected and subjected to further purification. EVs were isolated from the supernatant using differential ultracentrifugation, as described (Thery et al., 2006). Briefly, the supernatant was subjected to centrifugation at 10,000 × g for 30 min to remove large debris and unbroken cells. The resulting supernatant was then subjected to ultracentrifugation at 100,000 × g for 2 h to pellet the EVs. The EV pellet was resuspended in sterile phosphate-buffered saline (PBS) and washed by repeated centrifugation at 100,000 × g to remove residual contaminants. The purified EVs were stored at −80 °C until further use.

EVs were subjected to transmission electron microscopy (TEM) to determine EV morphology and approximate size. A sample of purified EVs was fixed with 2.5% glutaraldehyde and loaded onto carbon-coated copper grids. After staining with uranyl acetate, the grids were examined under a TEM (e.g., JEOL JEM-1400), allowing for high-resolution visualization of EVs’ shape, size, and distribution (van Niel et al., 2018). ImageJ software was used to measure the size of the vesicles from TEM images.

Additionally, nanoparticle tracking analysis (NTA) was employed to determine the size distribution and concentration of EVs in solution. EVs were diluted appropriately, and their motion was tracked using a NanoSight NS300 (Malvern Instruments), which provides size distributions in the range of 30–1,000 nm (Shao et al., 2018). This technique enabled the characterization of EVs’ concentration and particle size, which is critical for understanding their role in bacterial communication and immune modulation.

### Characterization of MAM-Containing EVs

Purified EVs were solubilized in SDS sample buffer and separated on a polyacrylamide gel. To confirm the presence of the MAM in purified EVs, the protein content was analyzed by Western blot. The gel was transferred to a nitrocellulose membrane, and the presence of the MAM protein was detected using anti-MAM polyclonal antibodies (U5893HC290_4: anti-CGVAPTKNTVKETEV (C-terminal, AA 118 – 131) and U5893HC290_8: anti-CTNVKGNPIEKNNFG (Loop, AA 88 – 101) from GenScript). Detection was performed using chemiluminescent substrates (e.g., SuperSignal West Pico Chemiluminescent Substrate, Thermo Fisher). For the identification and quantification of MAM in EVs, the MAM-containing EVs were also analyzed by antibodies specific to MAM protein. This allowed for a more quantitative assessment of MAM concentration in the EV preparations.

### MAM localization and Fractionation

To assess the localization of MAM in *F. prausnitzii* A2-165, a 30 mL culture of *F. prausnitzii* was pelleted by centrifugation, and the culture supernatant was concentrated to 2 mL. For the preparation of whole-cell lysate (WCL), the pellet was resuspended in 2 mL PBS and subjected to sonication (30 seconds on, 30 seconds off, 3 min total sonication time, at 70% amplitude using a cuphorn sonicator, and maintained at 4°C). The WCL was clarified by centrifugation at 20,800 x g for 15 min at 10°C. The resulting soluble fraction was designated as LS, while the insoluble fraction was resuspended in 50 µL PBS and designated as LP.

### Extracellular Vesicle (EV) Preparation

For EV preparation, 490 mL of *F. prausnitzii* culture was used per aliquot. The culture was concentrated down to 35 mL using a 10 kDa MWCO filter. Two conditions were prepared: one without proteinase K (PK) treatment, and one with PK treatment. In the PK treatment group, proteinase K was added to a final concentration of 100 µg/mL, and the mixture was incubated for 2 h at 50°C. Following PK treatment, phenylmethylsulfonyl fluoride (PMSF) was added to a final concentration of 5 mM to inhibit further proteolysis. The sample was not heat-killed to preserve vesicle structure and organization. The mixture was then centrifuged to harvest the EVs. The supernatants were collected for negative staining, and the pellets were resuspended in 20 mM Tris, 200 mM NaCl (pH 8.0).

### Electrophoresis of MAM Complexes

To assess the MAM oligomeric/complex composition, native PAGE was performed. The protein fraction was solubilized with 0.05% Dodecylmaltoside (DDM), which allowed for the separation of MAM oligomers/complexes in the 240-480 kDa range on a native gel. Negative staining was performed to confirm the impact of DDM on OMV integrity. Notably, 0.05% DDM was found to lyse OMVs, facilitating the analysis of the solubilized MAM complex.

### Statistical Analysis

Statistical analyses were performed using GraphPad Prism, and differences between treatment groups were analyzed using one-way ANOVA with Dunnett’s multiple comparisons. All results are presented as means ± standard error of the mean (SEM).

## Data availability

The authors declare that the data supporting the findings of this study are available within the paper and the Supplementary Information.

## Acknowledgments

We extend our gratitude to Elizabeth Skippington, Steven T. Rutherford, Sharookh B Kapadia, Man-Wah Tan for valuable insights and enlightening discussions. We thank Yamini Nanduri and David Castillo-Azofeifa for testing our supernatants and molecules in the intestinal regeneration assay. Our sincere appreciation goes to the teams at Genentech’s FACS and Luminex lab, Lab Animal Systems and Reports, Metabolomics lab for their expertise, technical guidance, and instrument support. Our special thanks to Saiyu Hang and Homer Pantua for their insight and assistance with the immune cell-based assays. We thank Tao Sun for generously providing the NF-κB NanoLuc reporter cell line (Invivogen) and the indispensable technical support. We thank Haoyuan Liu for his assistance in animal work. Figure panel in 1a was created using BioRender.

## Author contributions

L.Y., A.S., and D.W.W. conceptualized the study and designed the experiments. L.Y. conducted the *F. prausnitzii* in vitro assays. A.S. handled the purifications and characterizations of *F. prausnitzii* proteins and EVs. A.T.B. performed the electron microscopy analysis of *F. prausnitzii* EVs. H.Z. led the in vivo mouse studies. D.S. and Z.L. were responsible for metabolomics analysis from both in vitro bacterial supernatants. AC.L., J.K., and K.M.S. conducted the barrier function assays and analyses. E.K. provided bioinformatics support for genomic comparison analysis for *F. prausnitzii* isolates. L.Y., A.S., and D.W.W. wrote the manuscript, with contributions from all authors.

## Competing interests

L.Y., A.S., and D.W.W. declare no conflicts of interest. AC.L., A.T.B., H.Z., D.S., Z.L., J.K., and K.M.S. are employees of Genentech, Inc. and are Roche stockholders. Genentech, Inc./Roche did not influence the data collection, analysis, or the decision to submit the manuscript for publication. The authors affirm there is no financial gain nor patent applications pertaining to this publication to disclose. The authors affirm that the views and opinions expressed in this paper are those of the authors and do not necessarily reflect the views or interests of Genentech, Inc./Roche.

---

**Fig. S1 | Full panel of cytokine profiles from mouse sera in the DSS-colitis experiment.**

The full panel of cytokines and chemokines analyzed is shown. The following cytokines/ chemokines were also tested but were below the limit of detection: GM-CSF, IL-1β, IL-2, IL-3, IL-4, IL-10, IL-12 (p70), LIF, IL-15, MCP-1/CCL2, MIP-1α, MIP-1β. Statistical significance was determined using one-way ANOVA with Dunnett’s multiple comparisons.

**Fig. S2 | Correlation of the serum cytokines and body weight/ colon length from the *F. prausnitzii* DSS mouse experiment.**

Relationship between serum IL-17 (pg/mL) and body weight ratio (%) on different experimental days (**a**) and serum IL-17 (pg/mL) and colon length on the takedown day (**c**). Relationship between serum IL-6 (pg/mL) and body weight ratio (%) on different experimental days (**b**) and serum IL-6 (pg/mL) and colon length on the takedown day (**d**).

**Fig. S3 | Cell viability assays.**

Cell viabilities (**a**) for Fig. 2**a**, (**b**) for Fig. 2**c**, (**c**) for Fig. 2**d**, (**d**) for Fig. 2**e**, (**e**) for Fig. 2**f**, (**f**) for Fig. 3**c**, (**g**) for Fig. 4**a**, (**h**) for Fig. 4**b**, (**i**) for Fig. 5**c**, and (**j**) for Fig. 5**d**.

**Fig. S4 | Localization of MAM through fractionation of culture supernatants and proteinase K treatment and the function of EVs on T_H_17 cells.**

Silver stain (left) visualizes total proteins, while anti-MAM denaturing gels (right) detect MAM-specific signals with or without proteinase K treatment (**a**). Protein gels for different sample fractions (**b**).

**Fig. S5 | *F. prausnitzii* suppresses IL-8 secretion.**

Purified EVs from *F. prausnitzii* A2-165 supernatants showed no IL-8 inhibitory activity (**a**). However, whole supernatant, <10 kDa fraction of the supernatant, and the EV depleted supernatant of *F. prausnitzii* A2-165 culture effectively suppressed IL-8 secretion in HT-29 MTX cells when stimulated with 5 ng/mL of TNFα (**b**).

