## Supplemental for "The Secreted Metabolite Isopentenyladenine from *Faecalibacterium prausnitzii* Mitigates Gut Inflammation"

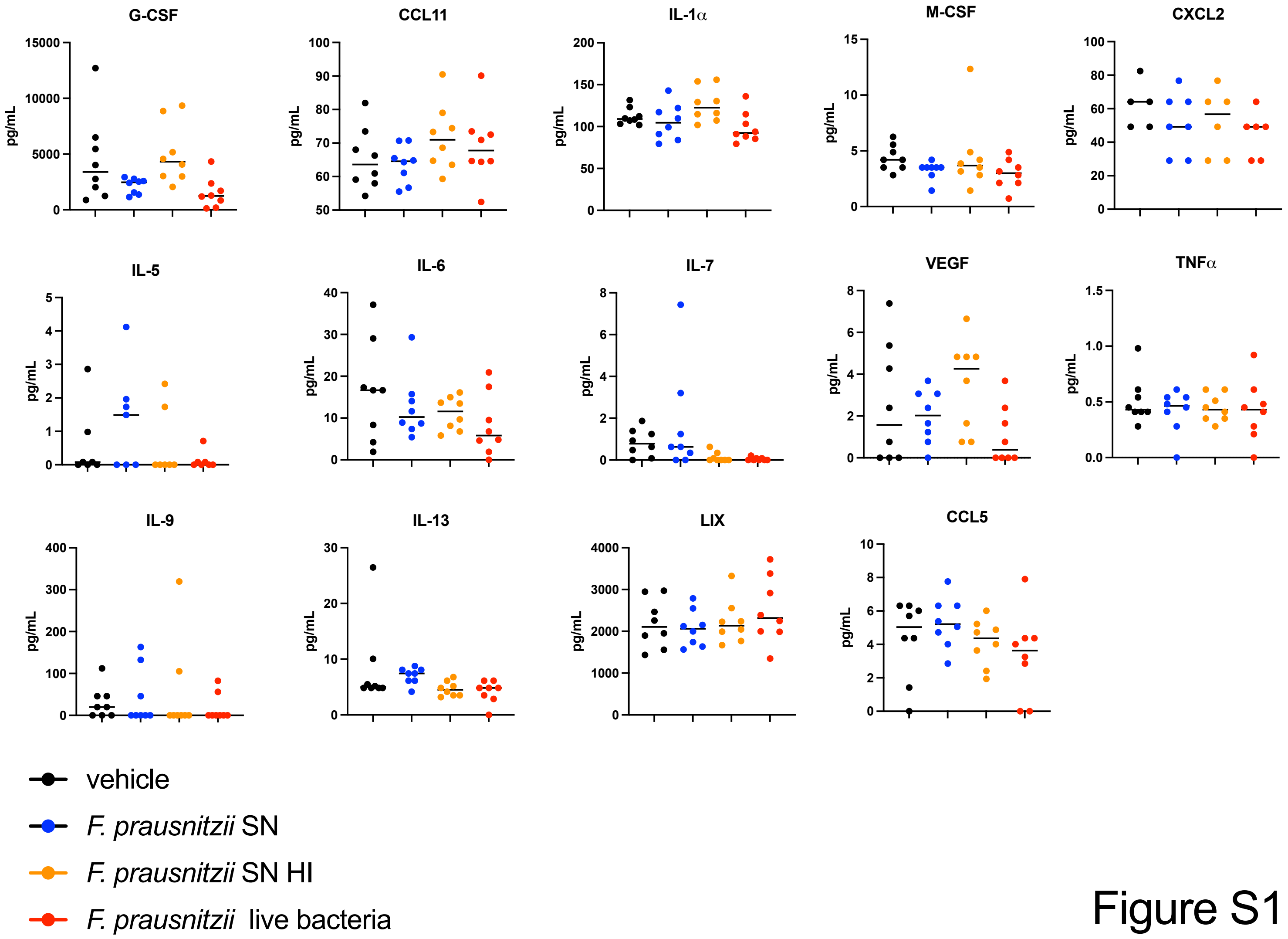
G-CSF
CCL11
IL-1α
M-CSF
CXCL2
IL-5
IL-6
IL-7
VEGF
TNFα
IL-9
IL-13
LIX
CCL5
pg/mL
vehicle
F. prausnitzii SN
F. prausnitzii SN HI
F. prausnitzii live bacteria

Figure S1

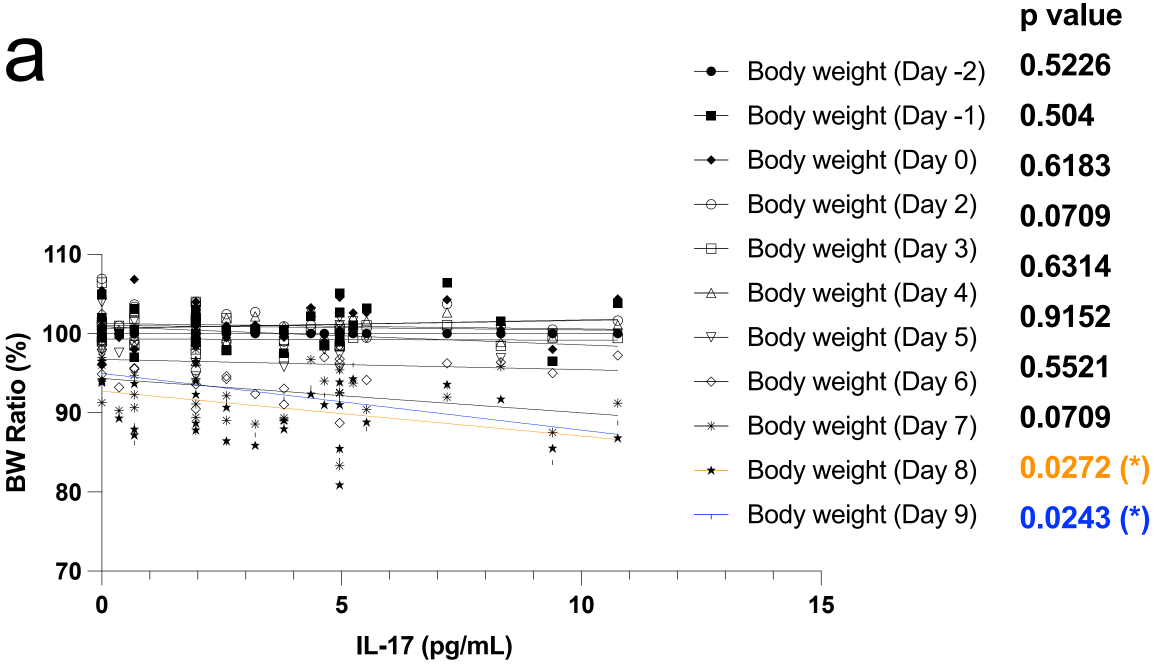

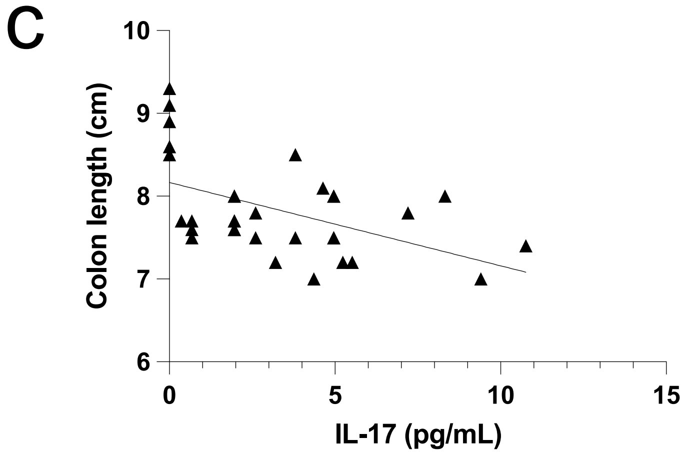

| | Colon length (cm) |
|---|---|
| P value | 0.0035 |
| Deviation from zero? | Significant |
| Equation | Y = -0.1006*X + 8.166 |

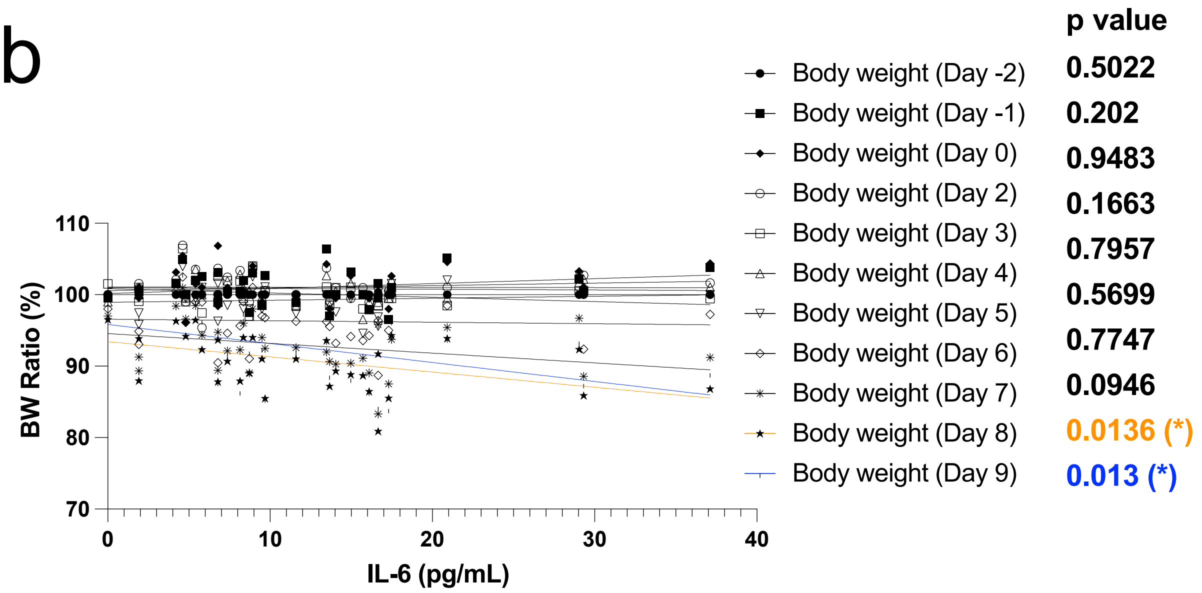

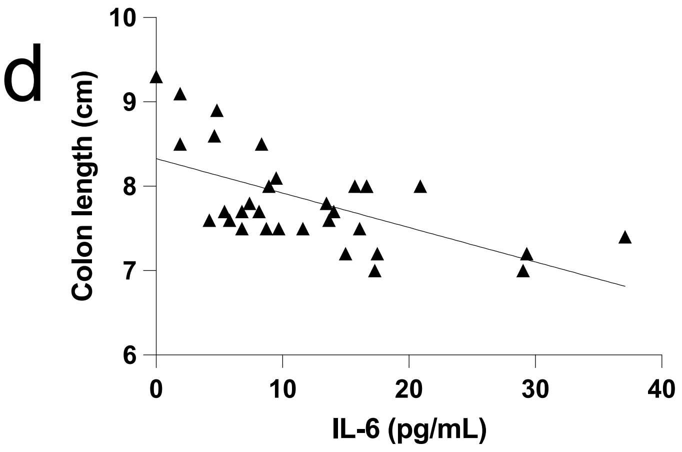

| | Colon length (cm) |
|---|---|
| P value | 0.0003 |
| Deviation from zero? | Significant |
| Equation | Y = -0.04081*X + 8.328 |

Figure S2

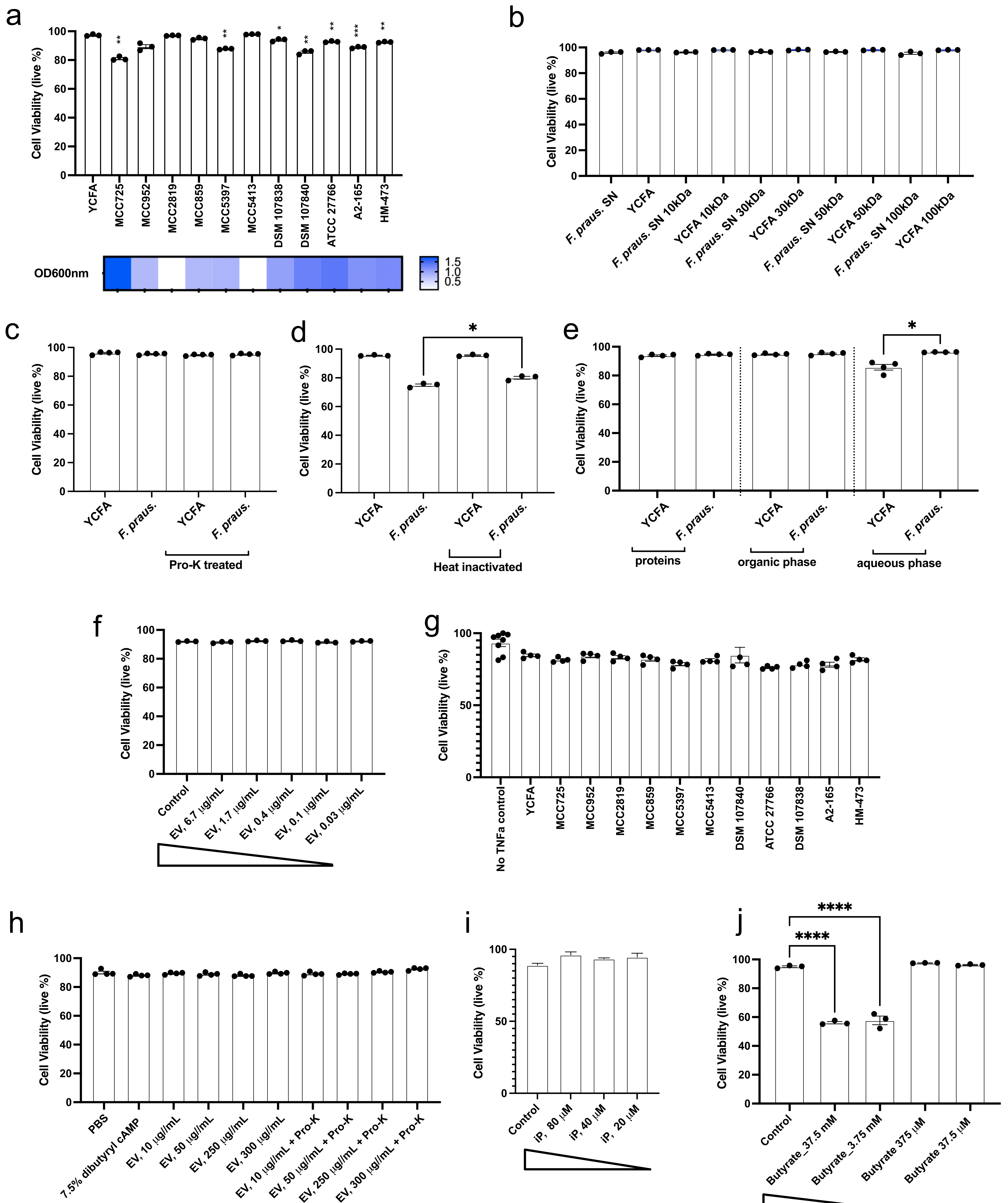
a
Cell Viability (live %)
YCFA
MCC725
MCC952
MCC2819
MCC859
MCC5397
MCC5413
DSM 107838
DSM 107840
ATCC 27766
A2-165
HM-473
OD600nm
b
F. praus. SN
YCFA
F. praus. SN 10kDa
YCFA 10kDa
F. praus. SN 30kDa
YCFA 30kDa
F. praus. SN 50kDa
YCFA 50kDa
F. praus. SN 100kDa
YCFA 100kDa
c
Pro-K treated
d
Heat inactivated
e
proteins
organic phase
aqueous phase
f
Control
EV, 6.7 μg/mL
EV, 1.7 μg/mL
EV, 0.4 μg/mL
EV, 0.1 μg/mL
EV, 0.03 μg/mL
g
No TNFa control
h
PBS
7.5% dibutyryl cAMP
EV, 10 μg//mL
EV, 50 μg//mL
EV, 250 μg//mL
EV, 300 μg//mL
EV, 10 μg//mL + Pro-K
EV, 50 μg//mL + Pro-K
EV, 250 μg//mL + Pro-K
EV, 300 μg//mL + Pro-K
i
iP, 80 μM
iP, 40 μM
iP, 20 μM
j
Butyrate_37.5 mM
Butyrate_3.75 mM
Butyrate 375 μM
Butyrate 37.5 μM

Figure S3

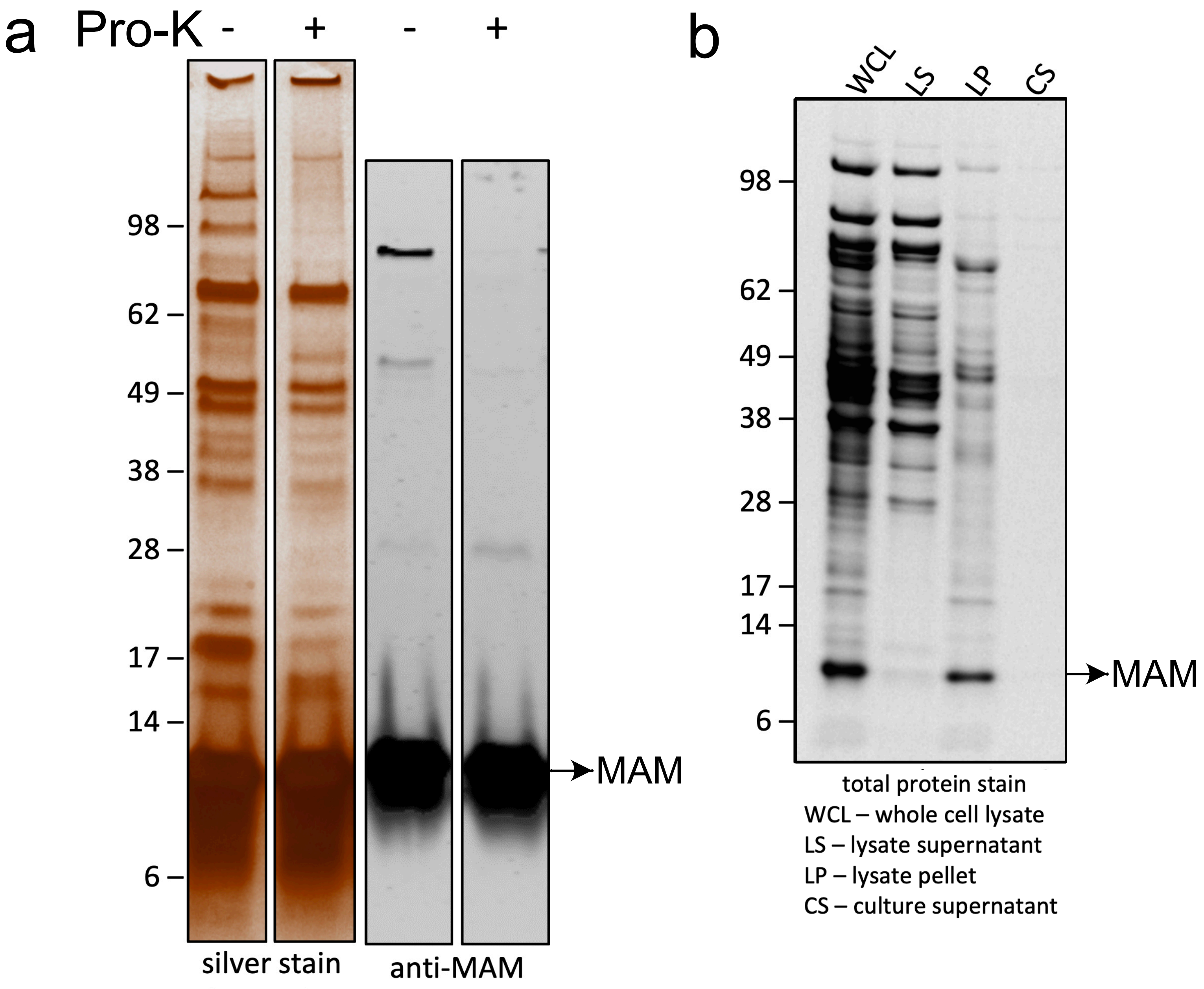
a
Pro-K - + - +
98
62
49
38
28
17
14
6
MAM
silver stain
denaturing
anti-MAM
denaturing
b
WCL
LS
LP
CS
98
62
49
38
28
17
14
6
MAM
total protein stain
WCL – whole cell lysate
LS – lysate supernatant
LP – lysate pellet
CS – culture supernatant

Figure S4

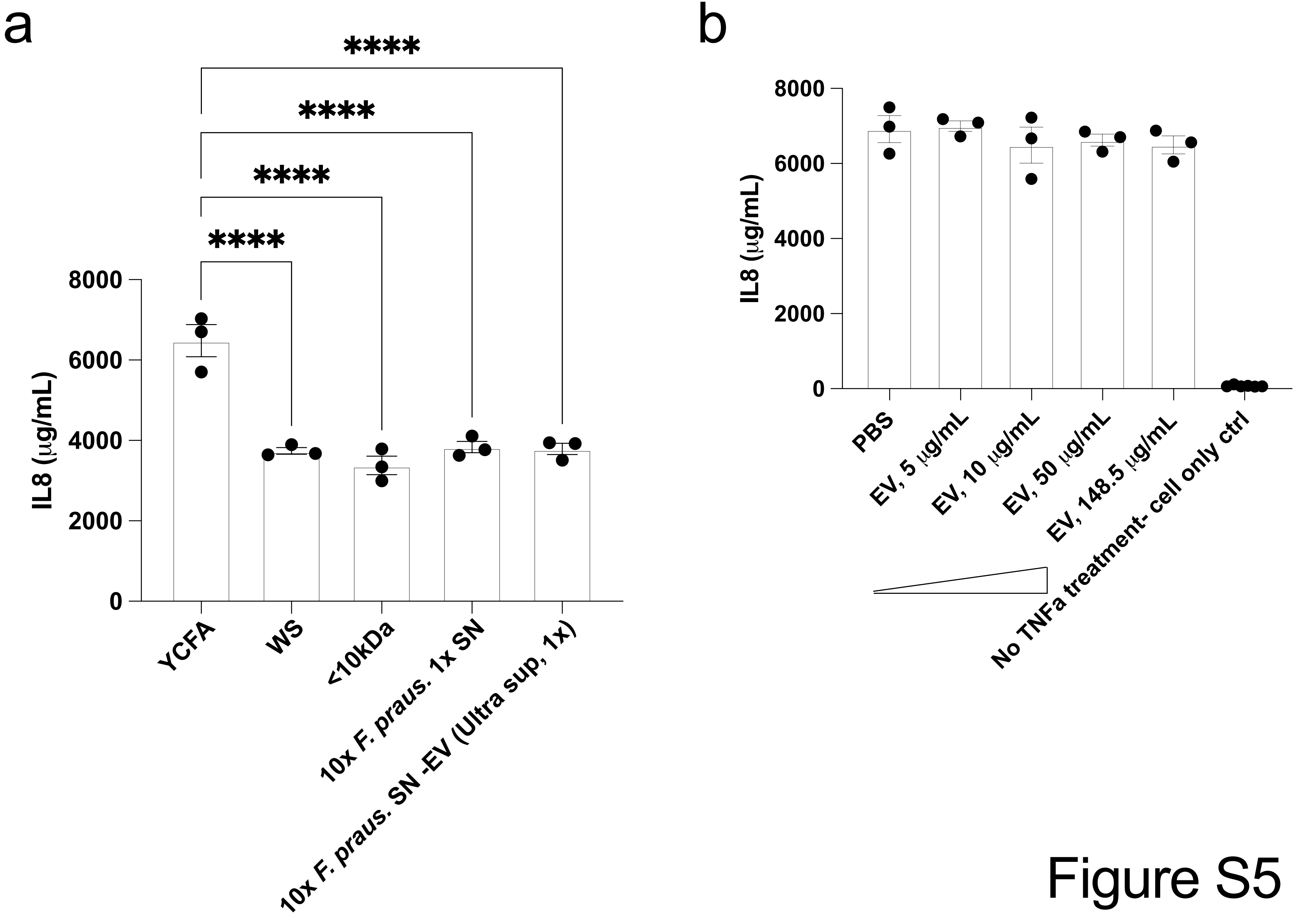
a
****
****
****
****
8000
6000
4000
2000
0
IL8 (μg/mL)
YCFA
WS
<10kDa
10x F. praus. 1x SN
10x F. praus. SN -EV (Ultra sup, 1x)
b
8000
6000
4000
2000
0
IL8 (μg/mL)
PBS
EV, 5 μg/mL
EV, 10 μg/mL
EV, 50 μg/mL
EV, 148.5 μg/mL
No TNFa treatment- cell only ctrl

Figure S5
